# Bright split red fluorescent proteins with enhanced complementation efficiency for the tagging of endogenous proteins and visualization of synapses

**DOI:** 10.1101/454041

**Authors:** Siyu Feng, Aruna Varshney, Doris Coto Villa, Cyrus Modavi, John Kohler, Fatima Farah, Nebat Ali, Joachim Dieter Mueller, Miri VanHoven, Bo Huang

## Abstract

Self-associating split fluorescent proteins (FPs) have been widely used for labeling proteins, scaffolding protein assembly and detecting cell-cell contacts. Newly developed self-associating split FPs, however, have suffered from suboptimal fluorescence signal. Here, by investigating the complementation process, we have demonstrated two approaches to improve split FPs: assistance through SpyTag/SpyCatcher interaction and directed evolution. The latter has yielded two split sfCherry3 variants with substantially enhanced overall brightness, facilitating the tagging of endogenous proteins by gene editing. Based on sfCherry3, we have further developed a new red-colored trans-synaptic marker called Neuroligin-1 sfCherry3 Linker Across Synaptic Partners (NLG-1 CLASP) for multiplexed visualization of neuronal synapses in living animals, demonstrating its broad applications.

## Introduction

Self-associating split fluorescent proteins (FPs) are a powerful tool for protein labeling and live-cell imaging. In this system, the eleventh β-strand of FP (FP_11_, 16 amino acids) is separated out from the remainder of FP (FP_1-10_) and is genetically fused to the protein of interest (POI). Specific fluorescence signal is detected when FP_1-10_ reconstitutes with the on-target FP_11_ to generate a functional fluorescent protein. Since the initial development of self-associating split GFP_1-10/11_ (*1*), this technology has been modified and adapted for an extensive range of applications including protein labeling and visualization (*2-4*), scaffolding protein assembly (*3*), protein solubility and aggregation assays (*5, 6*), and monitoring the membrane fusion process (*7*). One prominent application is generating a library of human cells with fluorescently tagged endogenous proteins via CRISPR/Cas9-mediated homology-directed repair (*4*). The small size of the GFP_11_ tag markedly improves the knock-in efficiency and simplifies the donor DNA preparation. In another application, the split GFP_1-10/11_ system has also been utilized to visualize synapses in living nervous systems by Neuroligin-1 GFP Reconstitution Across Synaptic Partners (NLG-1 GRASP) (*8*).

While all these applications have focused on a single, green-colored channel, expanding the color palette will greatly benefit the investigation of more complex biological systems by enabling multicolor imaging. For this purpose, we have recently developed a red-colored split sfCherry_1-10/11_ (*3*) and then a brighter split sfCherry2_1-10/11_ (*9*), enabling dual-color endogenous labeling in human cells using orthogonal FP_11_ tags (*9*) and visualization of *Listeria* protein secretion in infection (*10*). However, unlike split GFP_1-10/11_, which is as bright as its full-length, split sfCherry2_1–10/11_ produces substantially lower overall fluorescence signal than its full-length counterpart.

Here, we have characterized the complementation mechanism of split FP systems by examining their overall and single-molecule fluorescence brightness. The results suggest a 2-step complementation model in which the affinity between the FP_1-10_ and FP_11_ fragment is the major limitation to the overall fluorescence signal. Based on this model, we have devised a SpyTag/SpyCatcher-assisted approach to improve the complementation efficiency of sfCherry2_1-10/11_. Furthermore, we have engineered two split sfCherry3 variants with much-enhanced complementation efficiency through a combination of cycles of directed evolution and structure-based site-directed mutagenesis. For tagging endogenous proteins by gene editing, sfCherry3 improves the sorting efficiency for successfully knocked-in cells by 5-10 fold in six tested targets, as compared to sfCherry2. Moreover, we have also developed of a new red-colored trans-synaptic marker called Neuroligin-1 sfCherry3 Linker Across Synaptic Partners (NLG-1 CLASP). We established that like NLG-1 GRASP, NLG-1 CLASP labels connections between correct synaptic partners, and has a spatial pattern similar to that predicted by electron micrograph reconstruction (*11, 12*). As a validation, NLG-1 CLASP labeling is disrupted by loss of the *clr-1* gene, which is required for synaptic partner recognition (*13*).

## Results

### 1. The complementation efficiency of split sfCherry2 and split mNeonGreen2 can be explained by a dynamic association/dissociation equilibrium model

Previously, we have shown that split GFP_1-10/11_ has nearly identical overall brightness as its full-length counterpart, whereas both split mNeonGreen2_1-10/11_ (mNG2_1-10/11_) and split sfCherry2_1-10/11_ are substantially dimmer (*9*). This sub-optimal performance of mNG2_1-10/11_ and sfCherry2_1-10/11_ could be attributed to either (a) the lower molecular brightness of complemented split FPs or (b) incomplete complementation between the FP_1-10_ and FP_11_ fragments. To test the first possibility, we measured the single-molecule brightness of these three split FPs and their full-length counterparts in living cells using fluorescence fluctuation spectroscopy (*14*). We observed no significant difference in single-molecule brightness between the split and the full-length FPs in all three cases (Fig. 1A). Therefore, incomplete complementation should be the cause of the reduced overall fluorescence signal. The ratio of overall fluorescence between split and full-length FPs then reflects the complementation efficiency.

**Fig. 1.**
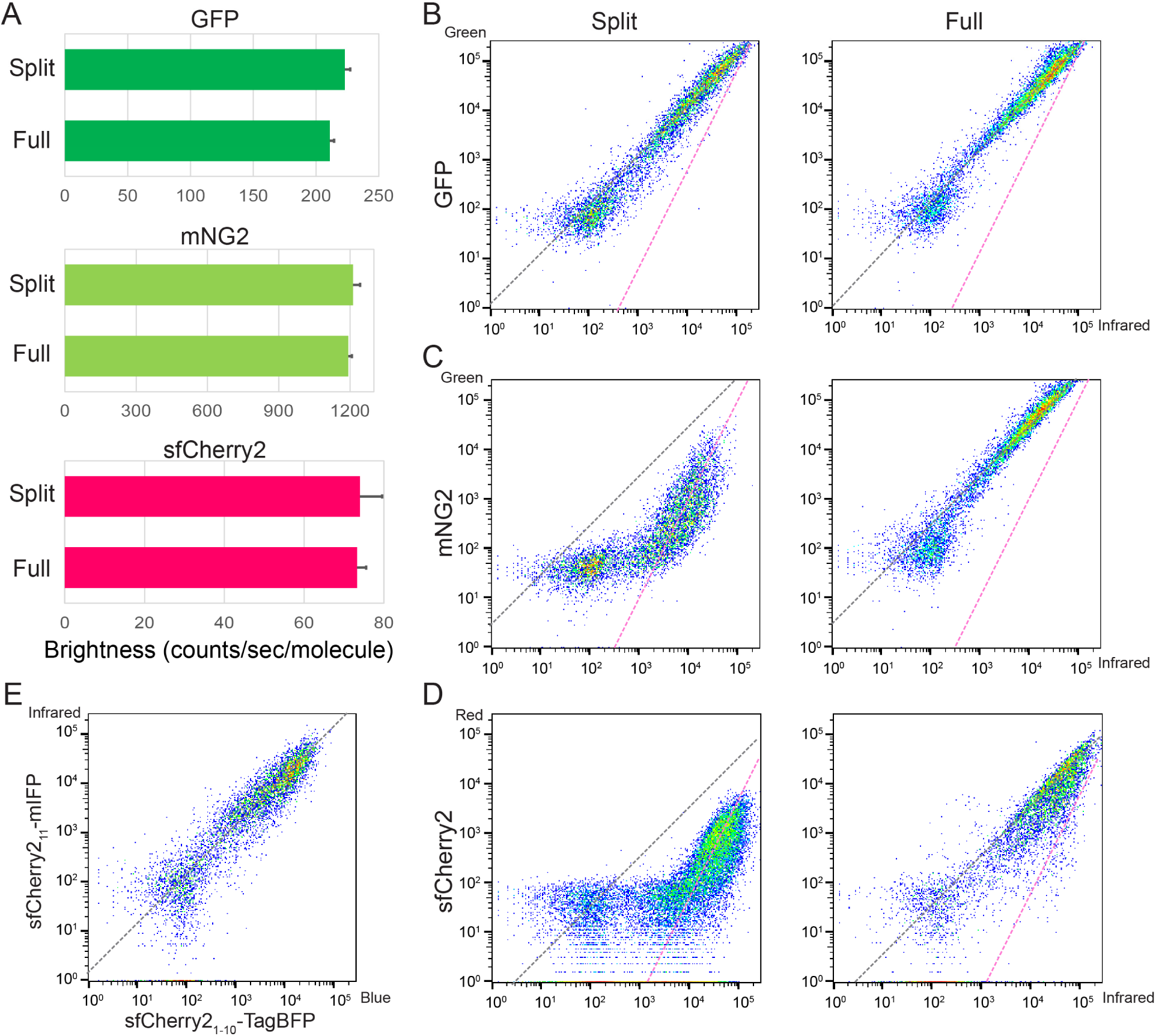
**Characterization of split fluorescent proteins**. **(A)** Single molecule brightness measurement of split FPs and their full-length counterparts using fluorescence fluctuation spectroscopy. *N* = 10. Error bars are standard deviations. **(B to D)** Flow cytometry analysis of whole cell fluorescence in HEK 293T expressing either **(B)** GFP_1-10/11_, **(C)** mNG2_1-10/11_, and **(D)** sfCherry2_1-10/11_ or their full-length counterparts. The x-axis is the log-scale infrared fluorescence intensity indicating the expression level, and the y-axis is the log-scale green (or red) fluorescence intensity. The grey dashed trend lines have a slope of 1 and intercepts are set to best follow the points in the right (full-length) panels. The pink dashed trend lines have a slope of 2 and intercepts set to best follow the points in the left (split) panels with the exception of GFP_1-10/11_. **(E)** Expression levels of two fragments are proportional within a wide range of expression levels in a co-transfect experiment. The grey dashed trend line has a slope of 1.

For mNG2_1-10/11_ and sfCherry2_1-10/11_, to determine their complementation efficiency and compare them to GFP_1-10/11_, we took a similar approach as previously done (*9*). We transiently expressed in HEK 293T cells the full-length FP or the two fragments: FP_1-10_ and FP_11_ on a well-folded carrier protein. We quantified whole cell fluorescence by flow cytometry while using a co-expressed infrared fluorescent protein mIFP to measure the expression level. mIFP was linked to the full-length FP or the FP_11_ fragment through a P2A self-cleavage site to ensure equimolar expression. Fluorescence intensities in infrared and green/red channels of each single cell events were displayed in log-log scale scatter plots (Figs. 1B-D).

In the cases of full-length FPs, as expected, single cell fluorescence intensities followed the trend line with a slope of 1 because mIFP was expressed equimolarly (Fig. 1E). In the split cases, split GFP_1-10/11_ almost completely followed the same diagonal line as its full-length counterpart (except for a subtle deviation at the lower-expression end) (Fig. 1B), suggesting a complementation efficiency of almost 100% across a wide range of expression levels. In contrast, both split mNG2_1-10/11_ and split sfCherry2_1-10/11_ deviated from the trend lines of their full length counterparts (Figs. 1C & D). This observation prompted us to consider a 2-step complementation process for self-associating split FPs: the two fragments undergo a reversible association/dissociation equilibrium before entering an irreversible process (fig. S1) of folding and/or chromophore maturation:

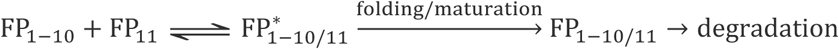

We have verified that in our co-transfection scheme, the expression levels (concentrations) of FP_1-10_ and FP_11_ fragments are proportional through an independent experiment in cells expressing mIFP-P2A-sfCherry2_11_-Carrier and TagBFP-P2A-sfCherry2_1-10_ (Fig. 1E). Then, at a steady state, this 2-step model predicts the following relationship between the two channels of flow cytometry (see Supplementary Notes):

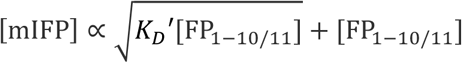

where [mIFP] represents the sum of concentrations of uncomplemented and complemented FP_11_, [FP_1–10/11_] is the concentration of matured FP_1-10/11_, and *K*_D_ ′ is the effective dissociation constant of the overall complementation/maturation process. When *K*_D_ ′ is much lower than the expression level of FP_11_ (the case of GFP_1-10/11_), the second term dominates, leading to a near proportional relationship between the two flow cytometry channels (slope of 1 in log-log plot, Fig. 1B). In the opposite case of high *K*_D_ ′ (the cases of mNG2_1-10/11_ and sfCherry2_1-10/11_), the first term dominates, resulting in log-log scatter plots following more closely to slope-2 trend lines, matching our observations (Figs. 1C&D).

### 2. SpyTag/SpyCatcher interaction improves complementation of split sfCherry2

Our model indicates that the complementation efficiency of split mNG2_1-10/11_ and sfCherry2_1-10/11_ improves with raised local concentration of fragments, leading to enhanced overall fluorescence signal. Therefore, we sought to utilize a pair of high-affinity binding partners to bring the two fragments into spatial proximity. Because a major advantage of the split FP_1-10/11_ is to label endogenous proteins through knocking-in the short FP_11_ peptide, it is preferable to have a small binding partner for the FP_11_ fragment. For this purpose, we chose the SpyTag/SpyCatcher (*15*) system, a peptide-protein pair that undergoes irreversible binding through formation of an isopeptide bond. The 13-amino-acid (aa) SpyTag is sufficiently short that even when concatenated with GFP_11_, the resulting sequence remains small enough for knock-in using synthetic oligo donor DNAs (*16*).

We examined SpyTag/SpyCatcher-assisted complementation on sfCherry2_1-10/11_ using the recently improved Spy002 pair (*17*) (Fig. 2A Upper panel). We fused SpyCatcher to the N-terminus of sfCherry2_1-10_ (the C-terminus is the split site) through a flexible linker in either 6 aa or 15 aa length (Fig. 2A Lower panel). We generated concatenated tags with SpyTag on either the N- or C-terminus of sfCherry2_11_ with a double-glycine linker. We performed similar flow cytometry experiments as described earlier except that mIFP was replaced by TagBFP. Among the four possible combinations of binders/tags (fig. S2), SpyCatcher-6aa-sfCherry2_1-10_ with SpyTag-sfCherry2_11_-TagBFP demonstrated the most pronounced shift towards a trend line with a slope of 1 in the scatter plot (Fig. 2B). We have further verified that a shorter SpyTag-sfCherry2_11_ fusion without the double-gylcine spacer behaved as efficiently (data not shown).

**Fig. 2.**
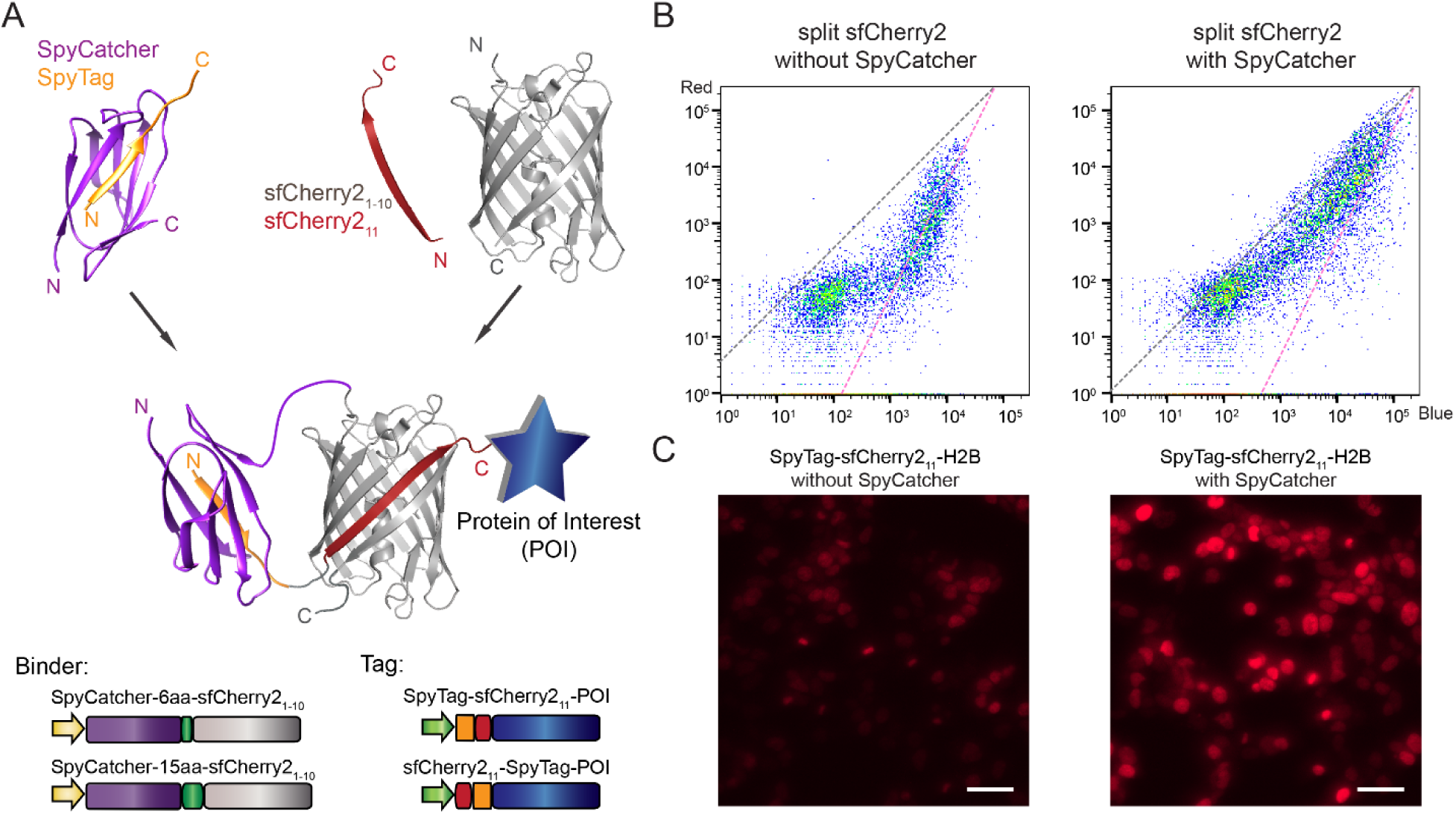
**SpyTag/SpyCatcher assisted complementation of split sfCherry2_1-10/11_**. **(A)** Schematic diagram of SpyTag/SpyCatcher-assisted complementation and construct design. **(B)** Flow cytometry analysis of whole cell fluorescence in HEK 293T cells expressing SpyTag-sfCherry2_11_-TagBFP with either sfCherry2_1-10_ alone or with SpyCatcher-6aa-sfCherry2_1-10_. The grey dashed trend line has a slope of 1 and the pink one has a slope of 2. **(C)** Fluorescence microscopy of sfCherry2_11_ labeled H2B without (left) or with (right) the assistance of SpyTag/SpyCatcher interaction. The imaging condition and brightness/contrast range were set the same for better comparison. Scale bars: 50 µm.

We validated the improvement in overall brightness for cellular microscopy by labeling the N-terminus of histone 2B (H2B) with SpyTag-sfCherry2_11_-TagBFP and co-expressing it with either sfCherry2_1-10_ or SpyCatcher-6aa-sfCherry2_1-10_ fusion in HEK 293T cells. We observed the Spy-assisted system (Fig. 2C) could mark the nuclei with much stronger fluorescence signal with the same expression vectors.

### 3. Engineering split sfCherry3 for better complementation efficiency

Previously, we engineered sfCherry2_1-10/11_ using a spacer-insertion strategy (*9*). This strategy was based on inserting a 32-aa spacer between the sfCherry_1-10_ and sfCherry_11_ coding regions. Beyond allowing mutagenesis of both fragments in a single PCR amplicon, the spatial constraints imposed by the linker were hypothesized to assist in the detection of the original mutations by raising the local concentrations of complimentary fragments. To increase the screening stringency for complementation-enhancing mutations, we chose to express the fragments of sfCherry2 as separate proteins using a pETDuet vector (Fig. 3A). Considering short peptides are prone to degradation in *E. coli*, we fused the sfCherry2_11_ sequence to the N-terminus of a well-folding carrier protein (SpyCatcher in our case).

**Fig. 3.**
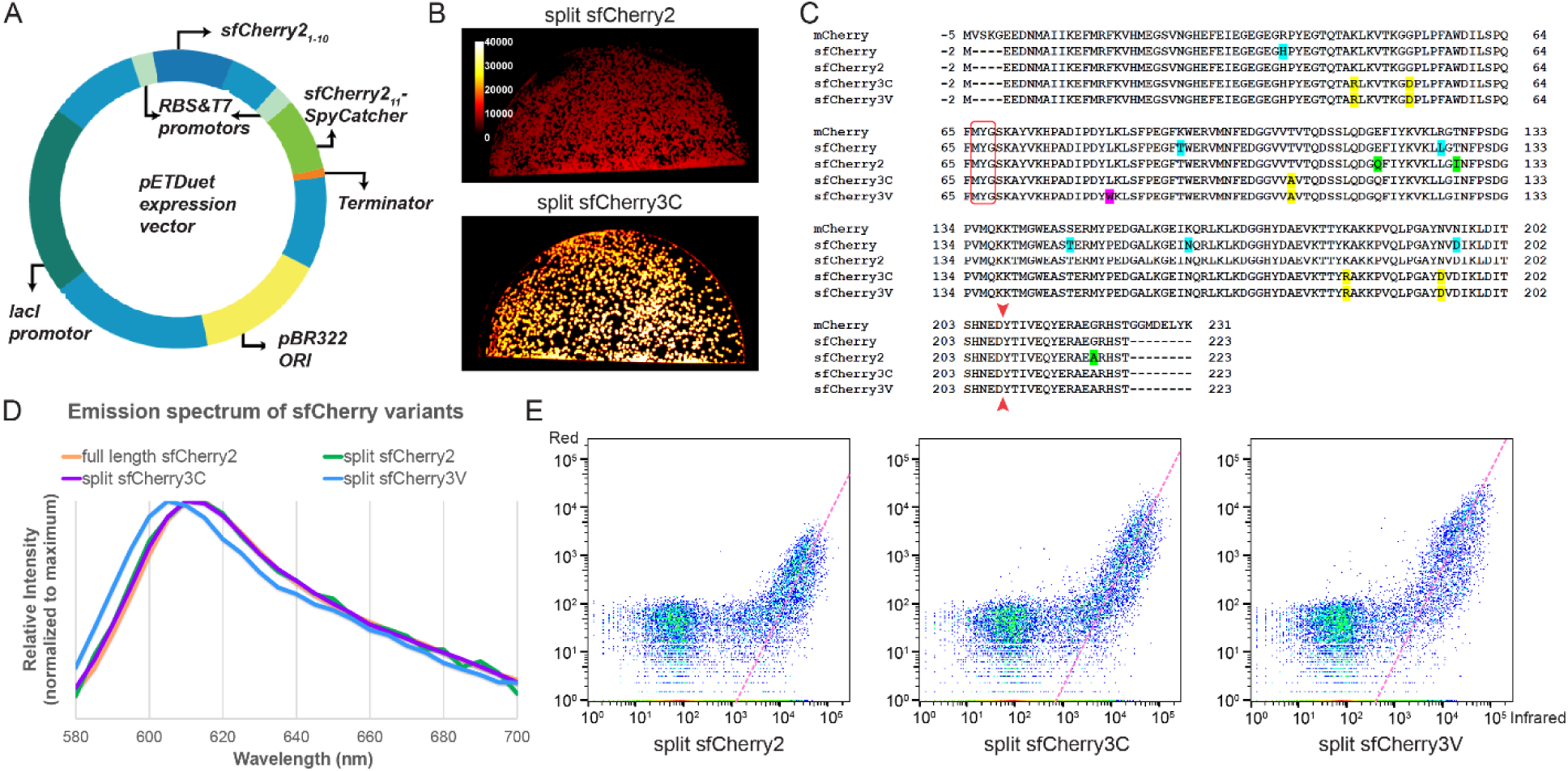
**Engineering and characterization of split sfCherry3**. **(A)** Schematics of pETDuet-based engineering platform. **(B)** Fluorescence images of E.coli colonies expressing split sfCherry2 or sfCherry3C from the pETDuet constructs. **(C)** Protein sequence alignment of mCherry, sfCherry, sfCherry2, sfCherry3C and sfCherry3V. The amino acids forming the chromophore are indicated by a red box. The split site is indicated by the red arrow. Starting from mCherry, mutations introduced in sfCherry, sfCherry2, sfChery3C and sfCherry3V are highlighted in cyan, green, yellow and magenta, respectively. The overall alignment numbering follows that of sfCherry. **(D)** Emission spectra of sfCherry variants. **(E)** Flow cytometry analysis of whole cell fluorescence in HEK 293T cells expressing mIFP-P2A-sfCherry2_11_-SpyCatcher and sfCherry2_1-10_ (left)/ sfCherry3C_1-10_ (middle)/ sfCherry3V_1-10_ (right). The pink dashed lines are results from linear least-square fitting with a fixed slope of 2 (see fig. S4).

We subjected the sfCherry2_1-10_ fragment to four rounds of error-prone PCR mutagenesis and screening. In every round, a mixture of ~20 brightest variants were selected for the next round. The final isolated mutants were then subjected to one round of DNA shuffling. We have not mutated sfCherry2_11_ so that all variants still bind the identical sfCherry2_11_ peptide. In the end, sfCherry3C with 5 substitutions, K45R, G52D, T106A, K182R, N194D (numbering starts from the 1^st^ Glu after the starting codon Met) was identified as the best variant after the directed evolution (Fig. 3B). All mutations were mapped to either surface orientated residues (T106A, K182R) or locations potentially interacting with the sfCherry2_11_ peptide (K45R, G52D, and N194D).

To further improve the complementation efficiency of sfCherry3C, we introduced rational mutations inspired by a mCherry mutant named cp193g7 that is tolerant of circular permutations near our split-site (*18*), because this mutant contains multiple similar mutations as in sfCherry (*19*) and sfCherry2 (*9*). We combinatorially introduced the remaining mutations of cp193g7 (I7F, F65L, and L83W) into sfCherry3C through site-directed mutagenesis. Only the variant containing a single L83W mutation gave brighter signal than sfCherry3C, which we designated as sfCherry3V (Fig. 3C). Complemented sfCherry3C_1-10/11_ has an identical emission spectrum as that of sfCherry2 (both split and full-length), whereas the emission spectrum of complemented sfCherry3V_1-10/11_ is blue-shifted by 5 nm (Fig. 3D). Fluorescence fluctuation spectroscopy indicates that sfCherry3C_1-10/11_ has the same single-molecule brightness as sfCherry2 (fig. S3). On the other hand, sfCherry3V_1-10/11_ is dimmer at the single-molecule level (fig. S3), which might be attributed to a difference in two-photon excitation of the blue-shifted chromophore at 1000 nm in the fluorescence fluctuation spectroscopy measurement.

Next, we performed the flow cytometry analysis (Fig. 3E) on HEK 293T cells co-transfected with mIFP-P2A-sfCherry2_11_-Carrier and sfCherryX_1-10_ (X being 2, 3C or 3V). By fitting the data points (excluding those below a threshold above the scattering background) to a line with a fixed slope of 2 (fig. S4, see Supplementary Notes), we found a substantial up-shift of the fitted lines from sfCherry2 to sfCherry3C and sfCherry3V. The shifts corresponded to increases in binding affinity by 2.5-fold and 8.2-fold, respectively, assuming the same single-molecule brightness under one-photon excitation for flow cytometry. This enhancement in binding affinity should lead to improved complementation efficiency (hence the overall brightness) at the same expression level of the FP_1-10_ fragment.

### 4. Efficient endogenous protein labeling in human cells using sfCherry3 variants

One unique application of FP_11_ tag is to generate library-scale fluorescently labelled endogenous proteins through genetic knock-in by homology-directed DNA repair. The small 16-aa size of FP_11_ allows us to fit its DNA sequence and short homology arms (~70 nt on either side) into commercially available 200 nt single-strand oligo-DNA (ssDNA). By electroporating Cas9/sgRNA ribonucleoprotein (RNP) and donor ssDNA into cells constitutively expressing the corresponding FP_1-10_ fragment, robust generation of FP-labeled human cell lines becomes fast and cost-effective (*4*). Multicolor knock-in has also been demonstrated by using orthogonal split FP systems to visualize differential distribution and interaction of multiple endoplasmic reticulum proteins (*9*).

The overall increased brightness of complemented sfCherry3 variants make them superior to sfCherry2 in the application of labeling endogenous proteins through knock-in. Because sfCherry3 and sfCherry2 share the same FP_11_ fragment, we adopted a reversed strategy as our previously reported one: knock-in of sfCherry2_11_ into HEK 293T wild-type cells through electroporation, followed by lentivirus infection for the three sfCherry_1-10_ variants (schematics in Fig. 4A). A total of six sfCherry2_11_ cell lines were created, with knock-ins at: lamin A/C (LMNA, inner nuclear membrane), clathrin light chain A (CLTA), RAB11A, heterochromatin protein 1 β (HP1b), endoplasmic reticulum proteins SEC61b (translocon complex) and ARL6IP1 (tubular ER). We compared Fluorescence-Activated Cell Sorting (FACS) enrichment efficiency for each cell line after infection with lentivirus (fig. 4B to 4G). In all examined targets, the sfCherry3C_1-10_ and sfCherry3V_1-10_ groups displayed remarkable population enhancement in the red-fluorescence-positive gate compared to sfCherry2_1-10_, rendering the sorting process substantially faster. Practically, for targets like CLTA, SEC61b or ARL6IP1, we were able to gate the fluorescent population around the clear peak and have 5-10 fold higher yield of isolating cells with successful knock-in.

**Fig. 4.**
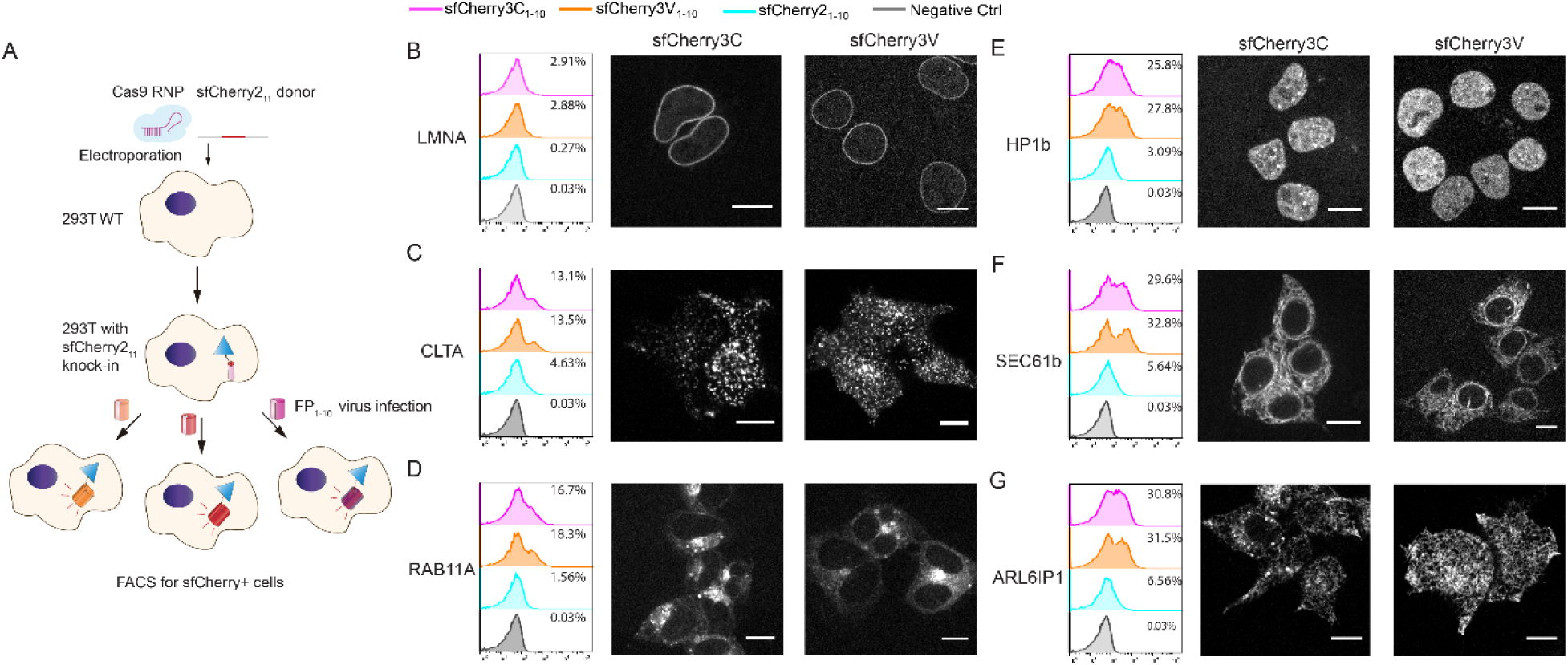
**Endogenous protein labeling in HEK 293T cells using sfCherry3 variants**. **(A)** Schematic diagram of knock-in followed by virus infection and FACS enrichment. **(B to G)** Analysis of FACS sorting efficiency in six targets, **(B)** lamin A/C, **(C)** clathrin light chain A, **(D)** RAB11A, **(E)** heterochromatin protein 1 β, **(F)** ER translocon complex SEC61b and **(G)** ER tubule protein ARL6IP1, and visualization of sorted knock-in cells through confocal fluorescence microscopy. Scale bar: 10 µm.

We further confirmed our knock-ins were on-target through confocal microscopy imaging (Fig. 4B to 4G). Noticeably, sfCherry3V_1-10_ groups have a reduced tendency to show fluorescent puncta from lysosomes (Fig. 4G) which is reported in our previous work (*9*). This makes the sfCherry3V_1-10_ the preferred protein fragment when labeling endogenous proteins involved in the endomembrane system.

### 5. NLG-1 CLASP visualizes specific subsets of synapses in live animals

To visualize synapses between specific sets of pre- and postsynaptic neurons in live animals, the trans-synaptic marker NLG-1 GRASP was designed using split GFP fragments (*8*). Complementary split GFP_1-10/11_ fragments were linked via a flexible linker to the transmembrane synaptic protein Neuroligin, which localizes to both pre- and postsynaptic sites in *C. elegans* (*8*). When the two neurons in which the complementary pre- and postsynaptic markers are expressed form synapses, the split GFP fragments come into contact, reconstitute and fluoresce (Fig. 5A). Using NLG-1 GRASP, we discovered that the recognition between two synaptic partners, the PHB sensory neurons and the AVA interneurons, is mediated the secreted ligand UNC-6/Netrin, its canonical receptor UNC-40/Deleted in Colorectal Cancer (*20*), and the receptor protein tyrosine phosphatase (RPTP) CLR-1 (*13*). NLG-1 GRASP has also been adapted to many other systems, indicating that this technology is transferable (*21*). The addition of a red fluorescent trans-synaptic marker would greatly expand this system.

**Fig. 5.**
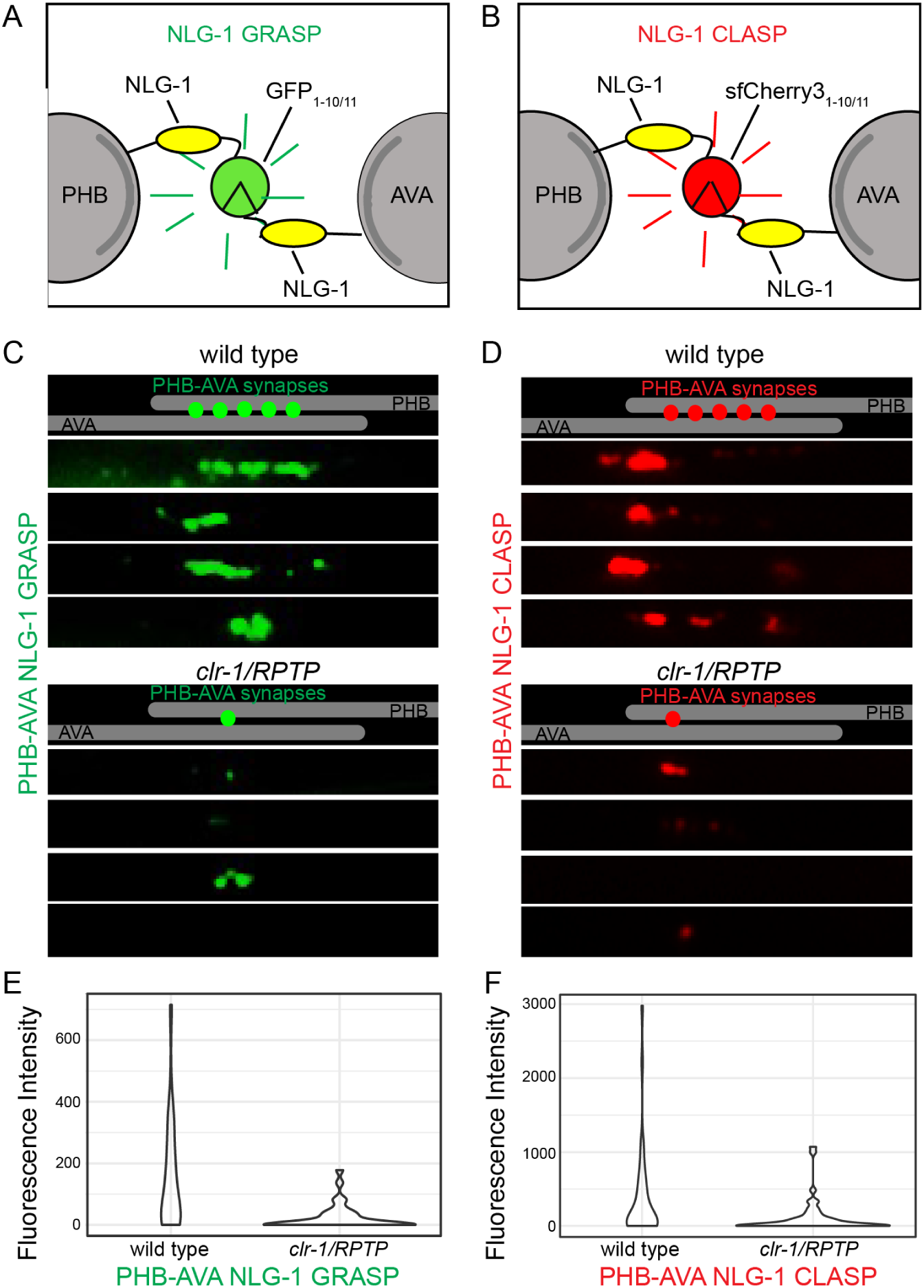
**NLG-1 CLASP visualizes specific subsets of synapses in live *C. elegans***. **(A & B)** Schematic diagram of GFP_1-10/11_-based NLG-1 GRASP and sfCherry3C_1-10/11_-based NLG-1 CLASP in PHB and AVA neurites. **(C & D)** Schematics and micrographs of NLG-1 GRASP and NLG-1 CLASP specifically labelling synaptic contacts between PHB and AVA neurons in *C. elegans*. PHB-AVA NLG-1 GRASP and NLG-1 CLASP fluorescence intensity are dramatically reduced in the synaptic partner recognition mutant *clr-1/RPTP*. **(E & F)** Quantification of the reduction in relative fluorescence intensities of NLG-1 GRASP and NLG-1 CLASP in *clr-1/RPTP* mutants. *N* ≥ 40.

Leveraging the brightened split-sfCherry3 tagging system, we have developed the split sfCherry-based NLG-1 CLASP. The left and right PHB sensory neurons, located in the posterior of *C. elegans*, form the majority of their synapses with the left and right AVA and PVC interneurons (*11, 12*). In this study, we focused on synapses formed between the left and right PHB neurons with the left and right AVA interneurons. To visualize PHB-AVA synapses with this newly developed split sfCherry3C, constructs were generated in which the sequence encoding the large fragment sfCherry3C_1-10_ was linked to the N-terminus of the *neuroligin-1 (nlg-1)* cDNA after the *nlg-1* signal sequence via a flexible 12GS linker. This half of the marker was expressed in PHB neurons using a promoter that within the posterior half of the worm, is specific for these presynaptic neurons (*_p_gpa-6*) (*22*). The complementary small fragment sfCherry3C_11_ was similarly linked to *nlg-1* and expressed in AVA neurons using a promoter that within the posterior half of the worm, is specific for these postsynaptic neurons (*_p_flp-18*) (*23*). Transgenic lines carrying both *_p_gpa-6::nlg-1::sfCherry3C_1-10_* and *_p_flp-18::nlg-1::sfCherry3C_11_* were generated. In wild-type animals, the distribution of red fluorescent puncta was similar to those observed in animals labeled with PHB-AVA NLG-1 GRASP (Fig. 5), and to that described by serial electron microscopy reconstruction (*11, 12*). To determine if these puncta were indeed synaptic, we introduced NLG-1 CLASP into *clr-1/RPTP* synaptogenesis mutants. In *clr-1/RPTP* mutants, PHB-AVA NLG-1 GRASP fluorescence intensity is dramatically reduced, and a PHB circuit-specific behavior is disrupted, indicating a reduction in synaptogenesis between the two neurons (*13*). We found that NLG-1 CLASP fluorescence intensity was also dramatically reduced in *clr-1/RPTP* mutants (Fig. 5), further indicating that NLG-1 CLASP puncta are synaptic.

## Discussion

Taken together, we have proposed a dynamic association/dissociation model to explain the complementation process of self-associating split FP system. We have performed whole cell fluorescence intensity measurement and single-molecule brightness quantification to validate this model. Inspired by our model, we have adopted the SpyTag/SpyCatcher system to enhance the local concentrations of fluorescent protein fragments, thereby improving its complementation efficiency and overall brightness. We have also designed an engineering strategy by expressing the two fragments individually through a pETDuet vector. Using this platform, we have developed a bright sfCherry3C_1-10/11_ (and its variant sfCherry3V_1-10/11_) with enhanced complementation efficiency, enabling the high-efficient generation of human cell lines with endogenously sfCherry-labeled proteins. Moreover, we have transformed the split sfCherry3C into a trans-synaptic marker called NLG-1 CLASP and have demonstrated the visualization of specific synapses in living animals.

The proposed 2-step model, consisting of a dynamic association/dissociation equilibrium followed by an irreversible folding/maturation process, can be generalized to other split fluorescent proteins, including the non-self-associating ones used to monitor protein-protein interaction in bimolecular fluorescence complementation (BiFC) assays (*24*) (*25*). In fact, there is no definitive boundary between the non-self-associating and self-associating split FPs. Instead, their only difference is in the spontaneous binding affinity, which can be characterized by *K*_D_′ in our model. For BiFC analysis, this affinity is one of the major determinant of the sensitivity. If it is too low, the split FP will fail to produce sufficient complementation signal even when the probed molecular interaction does occur. On the other hand, the opposite extreme leads to high background complementation as observed in certain split constructs (*6*). While this *K*_D_′ is not straightforward to measure biochemically *in vitro* due to the overall irreversible nature of complementation, our steady state flow cytometry analysis provides a reliable way to characterize it in cells. Our model also shows that the overall complemented signal is determined not only thermodynamically by the initial binding affinity and local fragment concentrations, but also kinetically by the rates of folding, chromophore maturation and protein degradation.

The assisted complementation demonstrated by SpyTag/SpyCatcher presents a simple way to improve complementation efficiency between FP fragments. This strategy can be expanded to multicolor imaging using FP_11_ tagging using orthogonal binder/tag pairs such as SnoopTag/SnoopCatcher (engineered by splitting an adhesin from *Streptococcus pneumoniae* (*26*)) or SsrA/SspB (a degradation tag and its adaptor protein from bacterial ClpX ATPase (*27, 28*)).

The substantial improvement of complementation efficiency in sfCherry3_1-10/11_ is beneficial for a wide variety of applications ranging from protein labeling to scaffolding protein complexes, as well as monitoring cell-cell connections. A highly efficient complementation process not only guarantees an enhanced overall brightness, but also enables us to tune down the expression level of FP_1-10_ fragments which might otherwise exhaust the cellular machinery that maintaining the protein homeostasis (biogenesis, folding, trafficking and degradation of proteins). This benefit is essential in scenarios that are sensitive to the expression of exogenous proteins, such as tagging endogenous proteins in embryos. Moreover, our engineering platform based on pETDuet vector can be utilized to optimize other self-associating split FPs with insufficient complementation efficiency, such as mNeonGreen2_1-10/11_.

Utilizing the split sfCherry3_1-10/11_ construct, we have established a completely orthogonal, red-colored trans-synaptic marker NLG-1-CLASP in addition to the original green fluorescent trans-synaptic marker NLG-1 GRASP. Similar cyan and yellow trans-synaptic markers (called dual-eGRASP) have recently been generated in vertebrates (*29*). The use of NLG-1 CLASP with these cyan and yellow trans-synaptic markers has the potential to allow simultaneous and differential labeling of synapses between a single neuron and three synaptic partners in live animals. Since many synaptic connections within the nematode have overlapping localizations within the nerve ring and other nerve bundles (*11, 12*), this tool will allow us to accurately visualize multiple subsets of a single neuron’s connections. These tools may similarly be of use in densely innervated regions of the vertebrate nervous system, such as the hippocampus and cortex. Thus, we propose that NLG-1 CLASP will be a powerful tool with which to probe the development and plasticity of neural circuits within live animals.

## Materials and Methods

### Molecular cloning

The DNA sequence of SpyCatcher 002 and SpyTag 002 (based on the reported sequence from (*17*)) were directly synthesized (Integrated DNA Technologies, IDT). The DNAs of histone H2B, TagBFP and mIFP were subcloned from mEmerald, TagBFP or mIFP fusion plasmids (cDNA source: the Michael Davidson Fluorescent Protein Collection at the UCSF Nikon Imaging Center) using Phusion High-Fidelity DNA Polymerase (Thermo Scientific). The P2A sequence are GCTACTAACTTCAGCCTGCTGAAGCAGGCTGGAGACGTGGAGGAGAACCCTGGACCT. The lentiviral plasmids pSFFV-GFP_1-10_, pSFFV-mNG2_1-10_ and pSFFV-sfCherry2_1-10_ were generated in our previous research (*9*). To build three pSFFV-mIFP-P2A-FP_11_-SpyCatcher constructs (used in Figs. 1A-C), three DNA fragments encoding mIFP, P2A-FP_11_ and SpyCatcher were ligated into linearized pSFFV vector (BamHI/NotI) using In-Fusion HD Cloning kit (Clontech) within one reaction. To construct pSFFV-TagBFP-P2A-sfCherry2_1-10_ plasmid (used in Fig. 1D), two DNA fragments encoding TagBFP or P2A were ligated in to linearized pSFFV-sfCherry2_1-10_ vector using In-Fusion. To generate pcDNA-SpyCatcher-6aa-sfCherry2_1-10_ and pcDNA-SpyCatcher-15aa-sfCherry2_1-10_ plasmids (used in Figs. 2A-B), PCR amplicons encoding SpyCatcher or sfCherry2_1-10_ were cloned into digested pcDNA3.1 vectors (HindIII/BamHI), and the different linkers were achieved through designing overlapping primers with various linker lengths. To make pSFFV-SpyTag-sfCherry2_11_-TagBFP and pSFFV-SpyTag-sfCherry2_11_-TagBFP-H2B (used in Figs. 2B-C), DNA fragments encoding SpyTag-sfCherry2_11_, TagBFP and H2B were ligated in to linearized pSFFV vector (BamHI/NotI) through In-Fusion.

The pETDuet-1 vector was kindly donated by Dr. Alexander Kintzer from Dr. Robert Stroud’s laboratory at UCSF. We generated the initial plasmid for mutagenesis screening by two rounds of In-Fusion ligation reaction: inserting the PCR amplicon encoding sfCherry2_1-10_ into the first multiple cloning site (MCS) of pETDuet-1 digested by NcoI, followed by inserting the DNA sequence encoding sfCherry2_11_-SpyCatcher into the second MCS of pETDuet-1 digested by NdeI. The NcoI restriction site preserved in the final product, but the NdeI restriction site was destroyed. For the mammalian expression and lentiviral production, DNAs of sfCherry3C_1-10_ were directly PCR amplified from identified pETDuet-1 construct (final mutant) and cloned into the lentiviral pSFFV vector (BamHI/NotI). To generate the sfCherry3V_1-10_ variant, we introduced the point mutation L83W into pSFFV-sfCherry3C_1-10_ plasmid using QuikChange II Site-Directed Mutagenesis Kit (Agilent Technologies). For the complete nucleotide sequence of sfCherry3C_1-10_, sfCherry3V_1-10_, SpyCatcher 002 and SpyTag 002, see Supplementary Table 1.

Constructs used in the NLG-1 CLASP application were generated using standard molecular techniques. To generate *_p_gpa-6::nlg-1::sfCherry3C_1-10_* construct (MVC227), *sfCherry3C_1-10_* was amplified from *pSFFV-sfCherry3C_1-10_* using the following primers: MVP846 (AGCTGCTAGCATGGAACGCATTTATCTTCTTCTCCTTCTTTTTCTGCCCAGGATACGATCCATGGAGGAGGACAACATGG) and MVP847 (TCCGGAGCTCGTCCTCGTTGTGGCTGGT). The fragment was subcloned into *_p_gpa-6::nlg-1::GFP_1-10_* (MVC6) (*20*), replacing GFP_1-10_, using the NheI and SacI sites. To generate the *_p_flp-18::nlg-1::sfCherry2_11_* (MVC228) construct, *sfCherry2_11_* was amplified from *H_2_B-sfCherry2_11_* using the following primers: MVP848 (AGCTGCTAGCATGGAACGCATTTATCTTCTTCTCCTTCTTTTTCTGCCCAGGATACGATCCTACACCATCGTGGAGCAGT) and MVP849 (TCCGGAGCTCGGTGCTGTGTCTGGCCTC). The fragment was subcloned into *_p_flp-18::nlg-1::GFP_11_* (MVC12)(*20*), replacing GFP_11_, using the NheI and SacI sites.

### Cell culture and lentiviral production

Human HEK 293T cells (UCSF cell culture facility) were maintained in Dulbecco’s modified Eagle medium with high glucose (Gibco), supplemented with 10% (vol/vol) FBS and 100 µg/ml penicillin/streptomycin (UCSF Cell Culture Facility). U2OS cells (American Type Culture Collection, Manassas, VA) were cultured in DMEM media, supplemented with 10% fetal bovine serum. All cells were grown at 37 °C and 5% CO_2_ in a humidified incubator. For the lentiviral production, 1 × 10^6^ HEK 293T cells were plated into T25 one day prior to transfection. 430 ng of pMD2.G plasmid, 3600 ng of pCMV-dR8.91 plasmid and 4100 ng of the lentiviral plasmid (pSFFV-sfCherry2_1-10_, pSFFV-sfCherry3C_1-10_ and pSFFV-sfCherry3V_1-10_) were co-transfected into HEK 293T cells using FuGENE HD (Promega) following the manufacturer’s recommended protocol. The virus containing supernatant is harvested 48 h after transfection and were centrifuged to pellet any packaging cells. Virus containing medium is used immediately or stored in −80 °C freezer for future use.

For single-molecule brightness measurement sample preparation, U2OS cells were grown in 24-well plates with #1.5 glass coverslip bottoms (In Vitro Scientific) and transfected ~24 hours before measurement using GenJet transfection reagent (SignaGen Laboratories) according to the manufacturer’s instructions. Immediately before measurement, the growth media was exchanged with PBS buffer with calcium and magnesium (Gibco).

### Sample preparation and data analysis in flow cytometry

To characterize the relationship between complementation efficiency and expression level in split GFP, split mNG2 and split sfCherry2, we made pSFFV-mIFP_P2A_full-length-FP and pSFFV-mIFP_P2A_FP_11_-SpyCatcher constructs. Corresponding to each scatter plots in Figures 1B-D, 3 × 10^4^ HEK 293T cells grown on 48-well plate (Eppendorf) were co-transfected with (B) left: 100 ng pSFFV-mIFP_P2A_GFP_11_-SpyCatcher with 200 ng pSFFV-GFP_1-10_, right: 100 ng pSFFV-mIFP_P2A_GFP[full-length] with 200 ng pSFFV-GFP_1-10_; (C) left: 100 ng pSFFV-mIFP_P2A_mNG2_11_-SpyCatcher with 200 ng pSFFV-mNG2_1-10_, right: 100ng pSFFV-mIFP_P2A_mNG2[full-length] with 200 ng pSFFV-mNG2_1-10_; (D) left: 100 ng pSFFV-mIFP_P2A_sfCherry2_11_-SpyCatcher with 200 ng pSFFV-sfCherry2_1-10_, right: 100 ng pSFFV-mIFP_P2A_sfCherry2[full-length] with 200 ng pSFFV-sfCherry2_1-10_. In Figure 1E, same cells were co-transfected with 100 ng pSFFV-mIFP_P2A_sfCherry2_11_-SpyCatcher and 200 ng pSFFV-sfCherry2_1-10_-TagBFP.

To test the Spy002 pair assisted complementation in sfCherry2, we built two tandem-binder constructs (pcDNA-SpyCatcher-6aa-sfCherry2_1-10_ and pcDNA-SpyCatcher-15aa-sfCherry2_1-10_) and two tandem-tag constructs (pSFFV-SpyTag-sfCherry2_11_-TagBFP and pSFFV-sfCherry2_11_-SpyTag-TagBFP). We then combinatorially co-transfected the same cells with 100ng tag construct plus 200 ng binder construct. In Figure 2B, to compare the complementation efficiency with or without Spy pair interaction, cells were co-transfected with either 100 ng pSFFV-SpyTag-sfCherry2_11_-TagBFP plus 200 ng pcDNA-sfCherry2_1-10_, or 100 ng pSFFV-SpyTag-sfCherry2_11_-TagBFP plus 200 ng SpyCatcher-6aa-sfCherry2_1-10_.

To validate the increased complementation of split sfCherry3 variants versus split sfCherry2, we made pSFFV-sfCherry3C_1-10_ and pSFFV-sfCherry3V_1-10_ constructs. Corresponding to each scatter plots in Figure 3E, same cells were co-transfected using with (left) 100ng pSFFV-mIFP_P2A_sfCherry2_11_-SpyCatcher with 200ng pSFFV-sfCherry2_1-10_; (middle) 100ng pSFFV-mIFP_P2A_sfCherry2_11_-SpyCatcher with 200ng pSFFV-sfCherry3C_1-10_; (right) 100ng pSFFV-mIFP_P2A_sfCherry2_11_-SpyCatcher with 200ng pSFFV-sfCherry3V_1-10_.

For flow cytometry analysis, 48 hours after transfection, transfected HEK 293T cells were digested with Trypsin-EDTA (0.25%) (Gibco) into single cells and re-suspended in 0.5 ml PBS solution. Analytical flow cytometry was carried out on a LSR II instrument (BD Biosciences) and cell sorting on a FACSAria II (BD Biosciences) in Laboratory for Cell Analysis at UCSF. Flow cytometry data analysis (gating and plotting) was conducted using the FlowJo software (FlowJo LLC).

### Single-molecule brightness measurement and data analysis

Fluorescence brightness measurements were carried out on a homebuilt two-photon microscope, which has been previously described (*30*). Pulsed laser light (100 fs pulses with a repetition frequency of 80 MHz) from a mode-locked Ti-Sapphire laser (Mai-Tai, Spectra Physics) was focused by a 63X C-Apochromat water immersion objective (NA = 1.2, Zeiss) to create two-photon excitation. The emitted fluorescence was collected by the same objective and separated from the excitation light by a dichroic mirror (675DCSXR, Chroma Technology). The fluorescence emission was separated into two detection channels by a 580nm dichroic mirror (585DCXR, Chroma Technology), and the green channel was further filtered by an 84nm-wide bandpass filter centered at 510 nm (FF01-510/84-25, Semrock). Avalanche photodiodes (SPCM-AQ-14, Perkin-Elmer) detected the fluorescence signal, and photon counts were recorded by a data acquisition card (FLEX02, Correlator.com) for ~60 seconds with a sampling frequency of 200 kHz. All measurements were carried out at an excitation wavelength of 1000 nm and a measured power after the objective of ~0.46 mW. The photon count record was analyzed to recover Mandel’s *Q* parameter as previously described (*31*) using programs written for IDL 8.7 (Research Systems, Inc.). The *Q* value is converted into brightness *λ*, which represents the average fluorescence intensity per molecule, using the relation *Q* = *γ*_2_*λT*, where *T* represents the sampling time and *γ*_2_ is a shape-dependent factor whose value has been determined as described previously (*32*).

### Mutagenesis and screening

The amino-acid sequence of sfCherry2 and the split site were from our previous published literature (*9*). The sfCherry2_1-10_ sequence was subjected to random mutagenic PCR (forward primer: AGGAGATATACCATGGAGGAGGACAAC, reverse primer: CTGCTGCCCATGTCAGTCCTCGTTGTG) using the GeneMorph II Random Mutagenesis Kit (Agilent Technologies). A high mutation rate protocol suggested in the instruction manual was adapted, with an initial target DNA amount of 0.2 µg and 30-cycle amplification. The cDNA library pool was gel-purified and ligated into a PCR-linearized pETDuet vector (forward primer: TGACATGGGCAGCAGCCA , reverse primer: CATGGTATATCTCCTTCTTAAAGTTAAACAAAATTATTTCTAGAGG. The product only contains the sfCherry2_11_-SpyCatcher coding sequence in the 2^nd^ MCS but not the sfCherry2_1-10_ in the 1^st^ MCS.) using In-Fusion. The plasmid pool was then transformed into *E.coli* BL21 (DE3) electrocompetent cells (Lucigen) by electroporation using the Gene Pulser Xcell™ Electroporation Systems (BioRad). The expression library was evenly plated on nitrocellulose membrane (Whatman, 0.45 µm pore size), which was sitting on an LB-agar plate with 30 mg/ml kanamycin. After overnight growth at 37 °C, the nitrocellulose membrane was carefully transferred onto a new LB-agar plate containing 1 mM Isopropyl-β-D-thiogalactoside (IPTG) and 30 mg/ml kanamycin and cultured for another 3-6 hours at 37 °C to induce the protein production. We performed the clone screening by imaging the IPTG-containing LB-agar plate using a BioSpectrum Imaging System (UVP). The brightest candidates in each library were pooled (typically ~20 from approximately 10,000 colonies) and served as templates for the next round of directed evolution. The DNA sequences of selected constructs were confirmed by sequencing (Quintara Biosciences). For DNA shuffling, we adopted the protocol described in Yu et al (*33*). Specifically, we PCR amplified the brightest six sfCherry3_1-10_ variants (forward primer: AGGAGATATACCATGGAGGAGGACAAC, reverse primer: CTGCTGCCCATGTCAGTCCTCGTTGTG) from the last round of random mutagenesis screening. PCR products of 651 bp were purified from 1% agarose gels using zymoclean gel DNA gel recovery kit (Zymo Research). The DNA concentrations were measured in Nanodrop and the fragments were mixed at equal amounts for a total of ~2 µg. The mixture was then digested with 0.5 unit DNase I (New England Biolabs) for 13min and terminated by heating at 95°C for 10 min. The DNase I digests were run on a 2% agarose gel, and the band with a size of 50-100 bp was selected and purified. 10 µl of purified fragments was added to 10 µl of Phusion High-Fidelity PCR Master Mix and reassembled with a PCR program of 30 cycles, with each cycle consisting of 95 °C for 60 s, 50 °C for 60 s and 72 °C for 30 s. After reassembly, 1 µl of this reaction was amplified by PCR. The shuffled library was then transformed, expressed and screened as described above. After the directed evolution was saturated (no apparent fluorescence increase in the induced colonies), the brightest clone was selected and the DNA sequences of the constructs were confirmed by sequencing (Quintara Biosciences). The emission spectra of split sfCherry variants (in *E.coli* solution culture expressing FPs) were measurement on a Synergy™ H4 Hybrid Multi-Mode Microplate Reader from UCSF Center for Advanced Technology.

### sgRNA production, RNP nucleofection and lentiviral transduction

We purchased synthetic single strand DNA oligos from Integrated DNA Technologies (IDT). We prepared sgRNAs and Cas9/sgRNA RNP complexes following our published methods (*4*). Specifically, sgRNAs were obtained by *in vitro* transcribing DNA templates containing a T7 promoter (TAATACGACTCACTATAG), an sgRNA scaffold region and the gene-specific 20 nt sgRNA sequence. DNA templates were produced by overlapping PCR using a set of 4 primers: 3 common primers (forward primer T25: 5’- TAA TAC GAC TCA CTA TAG -3’; reverse primer BS7: 5’- AAA AAA AGC ACC GAC TCG GTG C -3’ and reverse primer ML611: 5’- AAA AAA AGC ACC GAC TCG GTG CCA CTT TTT CAA GTT GAT AAC GGA CTA GCC TTA TTT AAA CTT GCT ATG CTG TTT CCA GCA TAG CTC TTA AAC -3’) and one gene-specific primer (forward primer 5’- TAA TAC GAC TCA CTA TAG NNN NNN NNN NNN NNN NNN NNG TTT AAG AGC TAT GCT GGA A -3’). For each template, a 50 μL PCR was performed with Phusion^®^ High-Fidelity PCR Master Mix (New England Biolabs) reagents with the addition of 1 μM T25, 1μM BS7, 20 nM ML611 and 20 nM gene-specific primer. The PCR product was purified and eluted in 12 μL of RNAse-free DNA buffer. Next, a 100-μL *in vitro* transcription reaction was performed with ~300 ng DNA template from PCR product and 1000 U of T7 RNA polymerase in buffer containing 40 mM Tris pH 7.9, 20 mM MgCl_2_, 5 mM DTT, 2 mM spermidine and 2 mM of each NTP (New England BioLabs). Following a 4 h incubation at 37°C, the sgRNA product was purified and eluted in 15 μL of RNAse-free RNA buffer. The sgRNA was quality-checked by running 5 pg of the product on Mini-PROTEAN TBE Precast Gels (Bio-Rad Laboratories) at 200-voltage for 60~80 min.

For the knock-in of sfCherry2_11_ into HEK 293T wild-type (WT) cells, 200-nt homology-directed recombination (HDR) templates were ordered in single-stranded DNA (ssDNA) from IDT. For the complete set of DNA sequence used for sgRNA in vitro transcription or HDR templates, see Supplementary Tables 2 and 3. Cas9 protein (pMJ915 construct, containing two nuclear localization sequences) was expressed in *E.coli* and purified by the University of California Berkeley Macrolab. HEK 293T WT cells were treated with 200 ng/mL nocodazole (Sigma) for ~17 hours before electroporation to increase HDR efficiency. 100 pmol Cas9 protein and 130 pmol sgRNA were assembled into Cas9/sgRNA RNP complexes just before nucleofection and combined with 150 pmol HDR template in a final volume of 10 μL. Electroporation was performed in an Amaxa 96-well shuttle Nuleofector device (Lonza) using SF-cell line reagents (Lonza). Nocodazole-treated cells were resuspended to 10^4^ cells/μL in SF solution immediately prior to electroporation. For each sample, 20 μL of cells was added to the 10 μL RNP/template mixture. Cells were immediately electroporated using the CM130 program and transferred to 48- well plate with pre-warmed medium. Electroporated cells were cultured and expanded for 7-10 days prior to lentiviral transduction.

One day before lentiviral transduction, knocked-in cells well split and seeded in 12-well plate at 6~8 × 10^4^ per well for 4 wells. The confluency should reach 70~80% on the day of transduction. The lentivirus titer of sfCherry2_1-10_, sfCherry3C_1-10_ or sfCherry3V_1-10_ was quantified independently by Lenti-X GoStix Plus kit (Takara, Cat# 631280) immediately before infection. And the supernatant was diluted (around 1:5 dilution) with fresh medium to make sure the final virus concentrations are the same across groups. The experimental groups were treated with 1ml diluted sfCherry2_1-10_, sfCherry3C_1-10_ or sfCherry3V_1-10_ viral supernatant supplemented with 10 ug polybrene (MilliporeSigma) respectively. The negative control was treated with fresh medium supplemented with the same concentration of polybrene. 24-hour after infection, the viral supernatant was swapped with fresh medium. After another 48 to 72 hours, the infected cells were harvested for flow cytometry analysis and cell sorting.

### Fluorescence Microscopy

To validate the Spy-assisted complementation can improve the overall brightness in protein labeling (Fig. 2C), we transfected HEK 293T cells (1.5 × 10^4^ per well) grown on an 8-well glass bottom chamber (Thermo Fisher Scientific) using FuGene HD (Promega). In order to achieve better cell attachment, 8-well chamber was coated with Fibronectin (Sigma-Aldrich) for 45 min before seeding cells. Total plasmid amount of 180 ng per well with the FP_11_ to FP_1-10_ ratio in 1:2 was used to achieve optimal labeling. 48 hours after transfection, cells were fixed with 4% paraformaldehyde and then imaged on a Nikon Ti-E inverted wide-field fluorescence microscope equipped with an LED light source (Excelitas X-Cite XLED1), a 40× 0.55 NA air objective (Nikon), a motorized stage (ASI) and an sCMOS camera (Tucsen).

Live-cell imaging (Figs. 4B-G) of sorted successful knock-in cells (with either sfCherry3C_1-10_ or sfCherry3V_1-10_ infection) was acquired on an inverted Nikon Ti-E microscope (UCSF Nikon Imaging Center), a Yokogawa CSU-W1 confocal scanner unit, a Plan Apo VC 100× 1.4 NA oil immersion objective, a stage incubator, an Andor Zyla 4.2 sCMOS camera and MicroManager2.0 software. Fluorescence recovery after photobleaching (FRAP) experiment (fig. S1) was performed on the same microscopy with a Vortran 473 nm laser for photobleaching and a Rapp Optoelectronic UGA-40 photobleaching system (COM5). The frame interval of the movie is 2.5 s and the total length is 10 min. The photobleaching was conducted by scanning the 473 nm (50 mW) laser over the region of interest at the 20 s of the movie. For optimal cell attachment, 8-well glass bottom chamber was coated with Fibronectin (Sigma-Aldrich) for 45 min before seeding sorted endogenously tagged HEK 293T cells. A background image (taken at the same imaging condition without putting on any sample) was subtracted from the live-cell microscopy images using the ImageJ software. Analysis of fluorescence microscopy images were performed in ImageJ.

### Generation of NLG-1 CLASP in *C. elegans*

All *C. elegans* strains were maintained using standard protocols(*34*) and were raised on 60 mm Nematode Growth Media plates seeded with OP50 *Escherichia coli* at 20 °C. Wild-type strains were *C. elegans* variety Bristol, strain N2, and the mutant strain used for this study was *clr-1(e1745) II*. Transgenic strains include *wyEx1982* (*20*), *wyEx1982; clr-1, iyEx368* and *iyEx368; clr-1. wyEx1982* contains extra-chromosomal PHB-AVA NLG-1 GRASP marker (*_p_gpa-6::nlg-1::GFP_1-10_* (60 ng/μl), *_p_flp-18::nlg-1::GFP_11_* (30 ng/μl), *_p_nlp-1::mCherry* (10 ng/μl), *_p_flp-18::mCherry* (5 ng/μl) and *_p_odr-1::DsRedII* (20 ng/μl)). *iyEx368* contains the extra-chromosomal PHB-AVA NLG-1 CLASP marker (*_p_gpa-6::nlg-1::sfCherry3C_1-10_* (68 ng/μl), *_p_flp-18::nlg-1::sfCherry2_11_* (38.6 ng/μl), *_p_nlp-1::GFP* (4.5ng/μl), and *_p_odr-1::DsRedII* (39 ng/μl)).

A Zeiss Axio Imager.A1 compound fluorescent microscope was used to capture images of live *C. elegans* under 63X magnification. Worms were paralyzed on 2% agarose pads using a 2:1 ratio of 0.3 M 2,3-butanedione monoxime (BDM) and 10 mM levamisole in M9 buffer. All micrographs taken of PHB-AVA NLG-1 GRASP and NLG-1 CLASP markers were of larval stage 4 animals. All data from micrographs were quantified using NIH ImageJ software. Intensity of PHB-AVA NLG-1 GRASP and PHB-AVA NLG-1 CLASP was measured as previously described (*13, 35*). Briefly, the intensity at each pixel within each synaptic puncta was measured using NIH ImageJ. To account for differences in background fluorescence, background intensity was estimated by calculating the minimum intensity value in a region immediately adjacent to the puncta. This minimum intensity value was then subtracted from the intensity for each pixel, and the sum of the adjusted values was calculated. For control, pictures of wild-type animals were also taken on the same day using the same settings.

## Acknowledgement

We thank Dr. David Alexander Brown, Dr. Yina Wang for extensive discussion on data analysis, Dr. Bin Yang for help in fluorescence light microscopy, Alejandro Ramirez for preparing reagents and supplies, Dr. Kari Herrington for help in FRAP experiment, Dr. Noelle L’Etoile for advice on NLG-1 CLASP experiments, Dr. Xiaokun Shu for sharing the BioSpectrum Imaging System, and Dr. Joseph DeRisi for generously sharing the nucleofector device. This work is supported by the National Institutes of Health R21EB022798 and R01GM124334 (to B.H.), UCSF Program for Breakthrough Biomedical Research (Byers Award in Basic Science to B.H.), the National Institutes of Health R01GM064589 (to J.K. and J.D.M.), the National Institutes of Health R01NS087544 and SC3GM089595 (to M.V.), T34GM008253 MARC fellowship (to N.A.), and the National Science Foundation 1355202 (to M.V.). B.H. is a Chan Zuckerberg Biohub investigator.

## Author Contributions

S.F. and B.H. conceived and designed the research; S.F. performed the flow cytometry, random mutagenesis and screening, CRISPR knock-in, FACS, confocal microscopy and FRAP experiments; J.K and J.D.M. performed single-molecule brightness measurement and analysis; S.F. and C.M. performed the molecular cloning and site-directed mutagenesis experiments; A.V., D.C.V., F.F., N.A. and M. V. perfored the NLG-1 CLASP experiments and analysis; S.F. and B.H. analyzed the data; S.F. and B.H. wrote the manuscript.

## Supplementary Figures

**Supplementary Figure 1.**
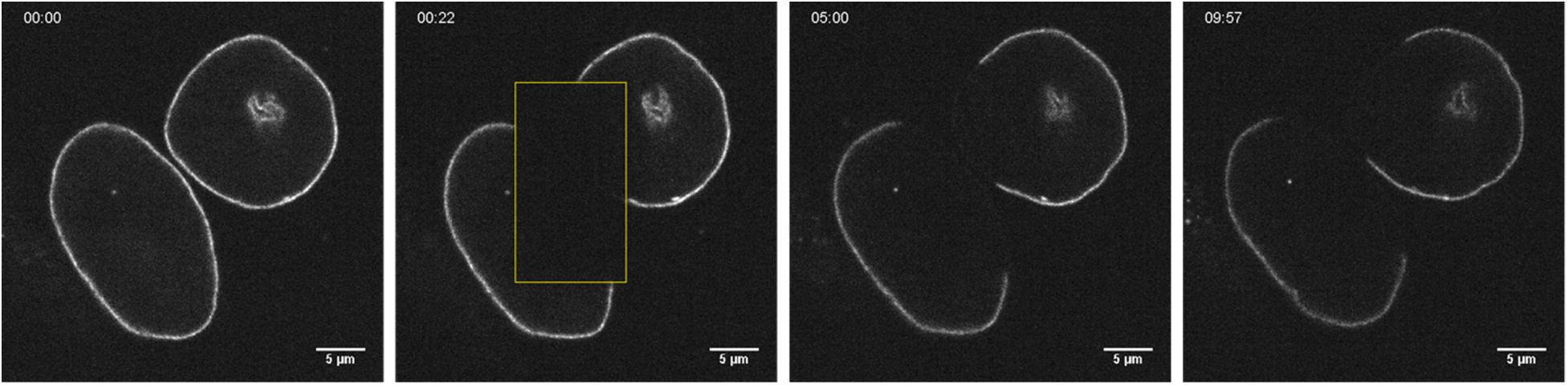
Snapshots of fluorescence recovery after photobleaching (FRAP) experiment on endogenously mNeonGreen2_1-10/11_ labeled Lamin A/C (nuclear envelope) in HEK 293T cells at different time points (before bleaching, 1^st^ frame, 5 min and 10 min after bleaching). The yellow square marks the photobleaching area.

**Supplementary Figure 2.**
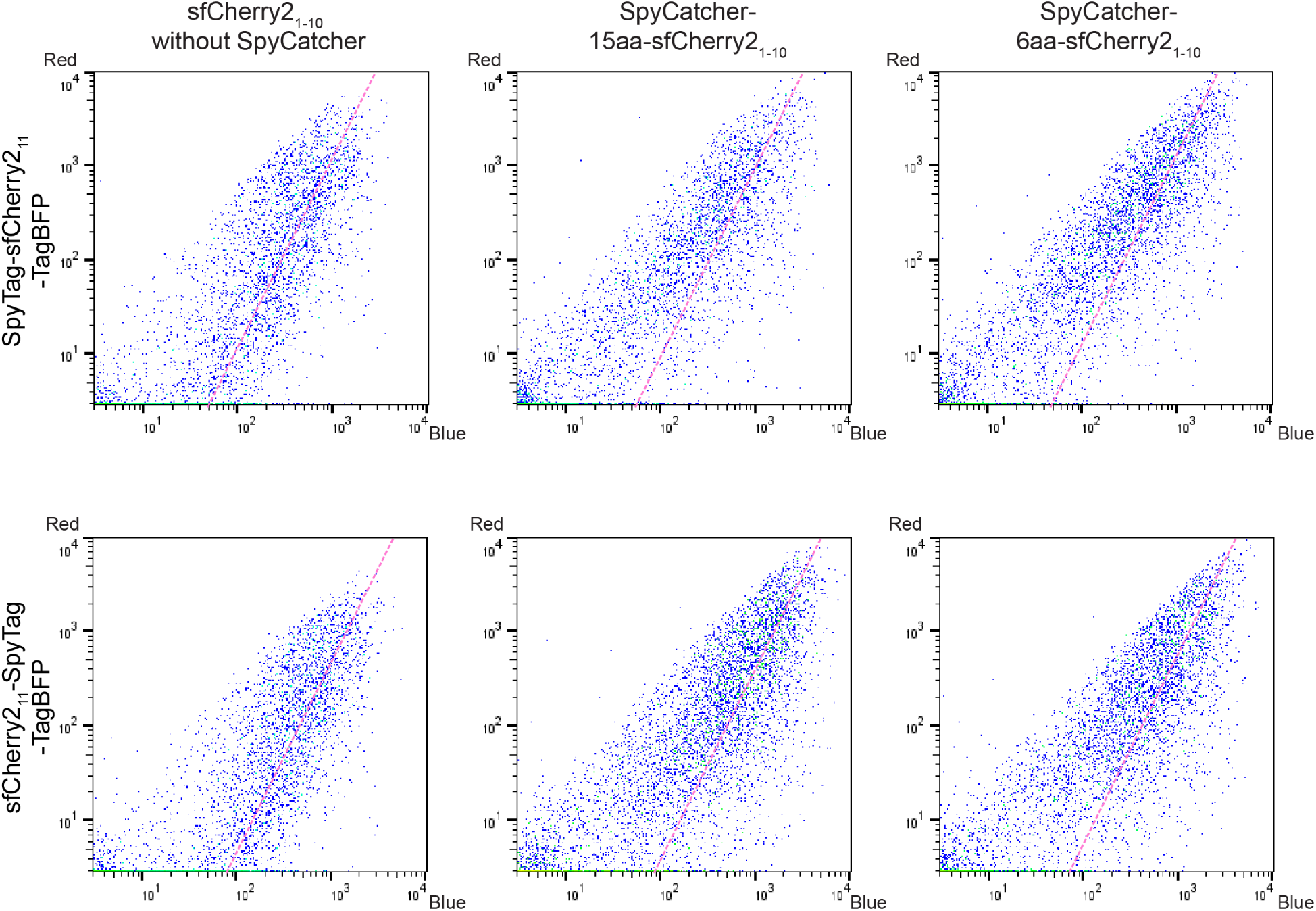
Flow cytometry analysis of whole cell fluorescence in HEK 293T cells expressing different tandem tags (upper: SpyTag-sfCherry2_11_-TagBFP, lower: sfCherry2_11_-SpyTag-TagBFP) with different binders (left: sfCherry2_1-10_ alone, middle: SpyCatcher-15aa-sfCherry2_1-10_, right: SpyCatcher-6aa-sfCherry2_1-10_). The pink dashed trend line has a slope of 2.

**Supplementary Figure 3.**
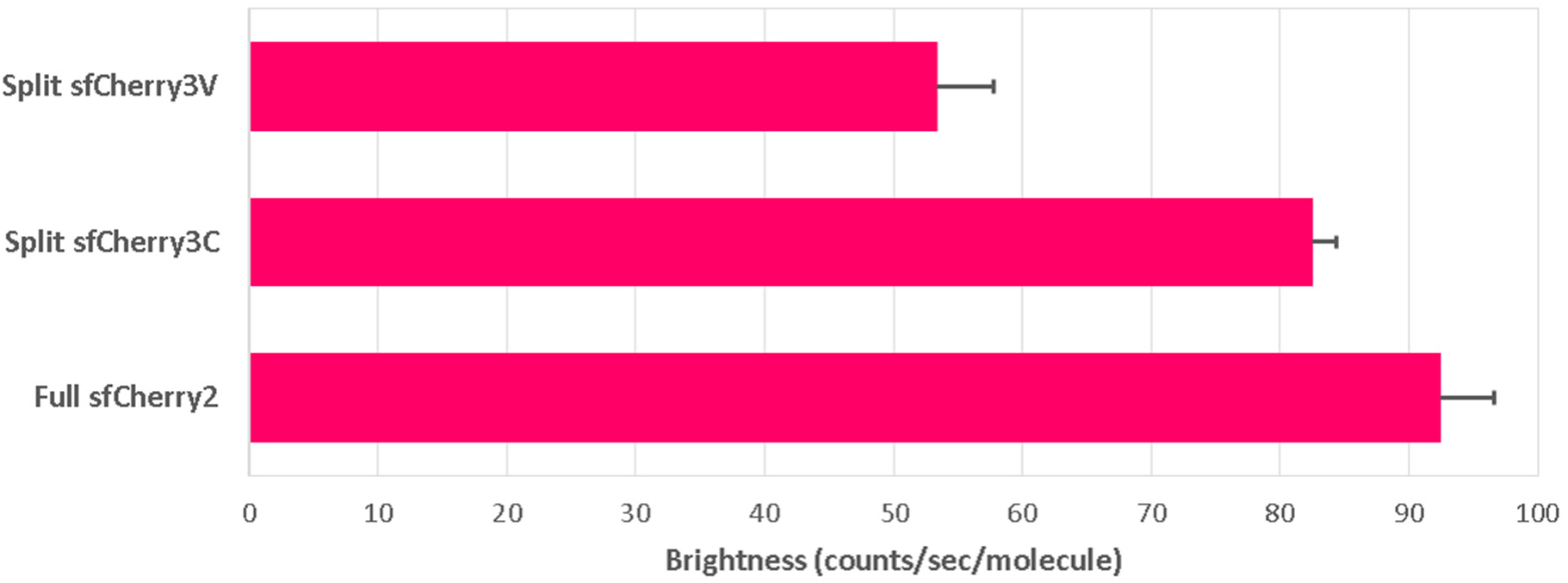
Single-molecule brightness measurement of split sfCherry3V, split sfCherry3Cs and full-length sfCherry2 using fluorescence fluctuation spectroscopy by two-photon excitation.

**Supplementary Figure 4.**
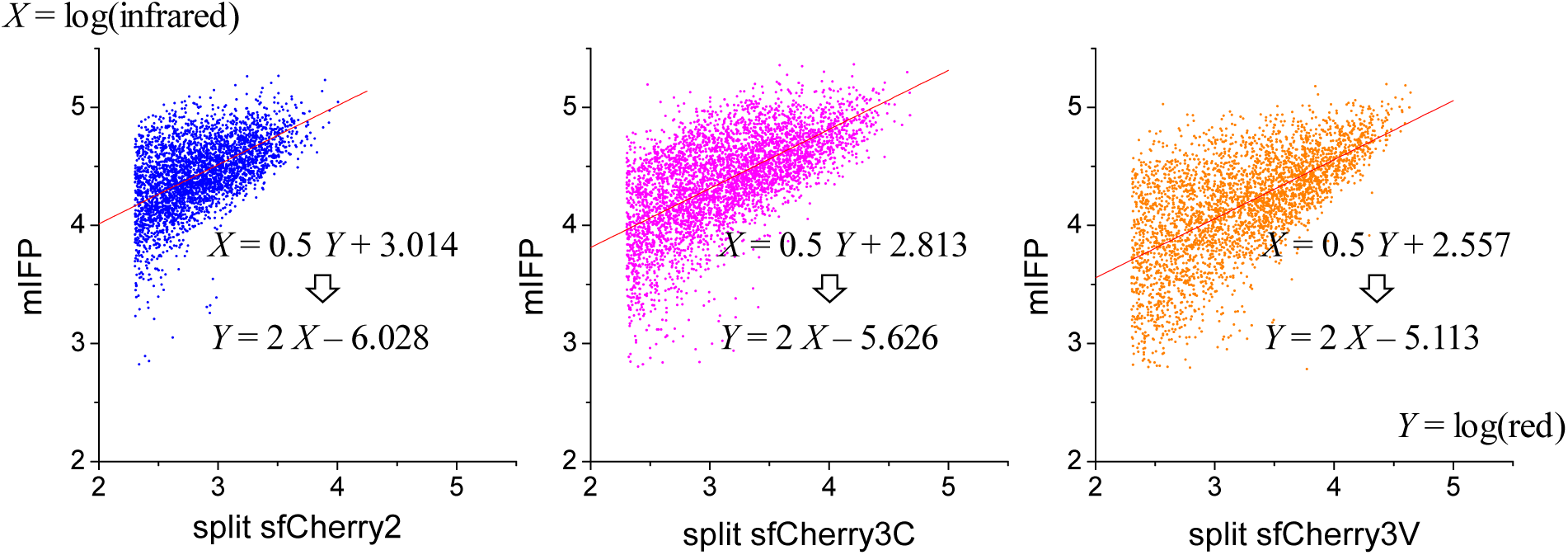
Least square linear fitting of flow cytometry points in Figure 3E with a fixed slope of 0.5. The *x*- and *y*-axes were flipped compared to main figures in order to properly handle the asymmetric errors in the two channels. Points with intensities within 3σ of the scattering background in either channel were excluded from the fitting. The intercept change in the fitted line (from 3.014 ± 0.004 to 2.813 ± 0.005 to 2.557 ± 0.006, ± standard error for least square fitting) was used to calculate the relative change in dissociation constant *K_D_*′ (see Supplementary Notes).

## Supplementary Notes

### A two-step model for FP_1-10/11_ complementation

The overall complementation process of commonly used split fluorescent proteins is known to be irreversible [1, 2]. Here, we verified that in the case of the self-complementing split mNG2_1-10/11_, the overall process is also irreversible by fluorescence recovery after photobleaching (FRAP) experiment. In HEK 293T cells with lamin A/C endogenously labeled by mNG2_11_ and constitutively overexpressed mNG2_1-10_ [3], bleaching a part of the nuclear lamina led to no observed fluorescence recovery in 10 min (Supplementary Figure 1, Supplementary Movie). Considering the fast maturation of mNeonGreen (< 10 min) [4], this lack of fluorescence recovery indicated that there was no exchange between the complemented mNG2_1-10_ with free cytosolic mNG2_1-10_.

On the other hand, a one-step, irreversible complementation would result in a direct proportional relationship between the complemented fluorescence signal and the fragment expression level. In this case, the FACS data points in Figure 1C (left) should follow a trend line with a slope of 1, whereas in reality, they fall closer to a trend line with a slope of 2. To explain this observation, we consider a two-step complementation model, in which the first step is a relatively fast reversible binding between the FP_1-10_ and FP_11_ fragments, forming an intermediate FP^*^_1-10/11_ complex, followed by a second, slower step of irreversible folding and/or fluorophore maturation.

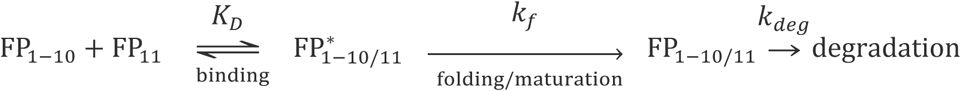

At a steady state, the concentration of the complemented, functionally fluorescent species, FP_1-10/11_, is related to the concentration of the intermediate complex, FP^*^_1–10/11_ by:

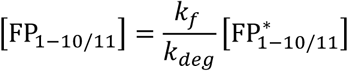

where *k_f_* and *k_deg_* are the folding/maturation rate and degradation rate, respectively. Generally, we would expect that the folding rate is much faster than the degradation rate, i.e. *k_f_* >> *k_deg_* and hence [FP_1–10/11_] >> [FP^∗^_1–10/11_]. The concentration of FP^*^_1-10/11_ is further connected to the concentration of the free fragments as:

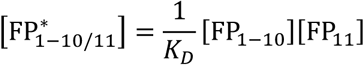

where *K_D_* is the dissociation constant for the initial binding step. In our experiment, FP_11_ and FP_1-10_ are expressed from co-transfected plasmids. Figure 1E indicates that their expression levels in individual cells follow a proportional relationship:

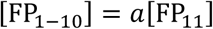

where *a* should be close to 1 considering that the same expression vectors are used for FP_11_ and FP_1-10_. Taken together, we have:

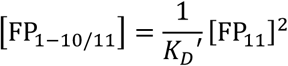

where the effective dissociation constant

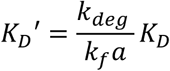

In our FACS experiment, the complemented fluorescence signal, *Y*, is proportional to [FP_1–10/11_], whereas the expression marker (mIFP) signal, *X*, is actually proportional to the total concentration of complemented and uncomplemented FP_11_ fragment

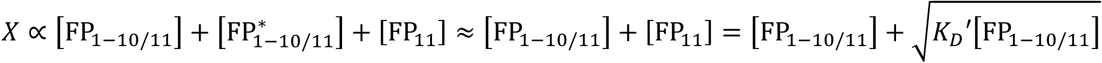

considering that [FP_1–10/11_] >> [FP^∗^_1–10/11_].

The relationship between *Y* and *X*, on a log-log plot, follows the black curve on the simulated plot below:

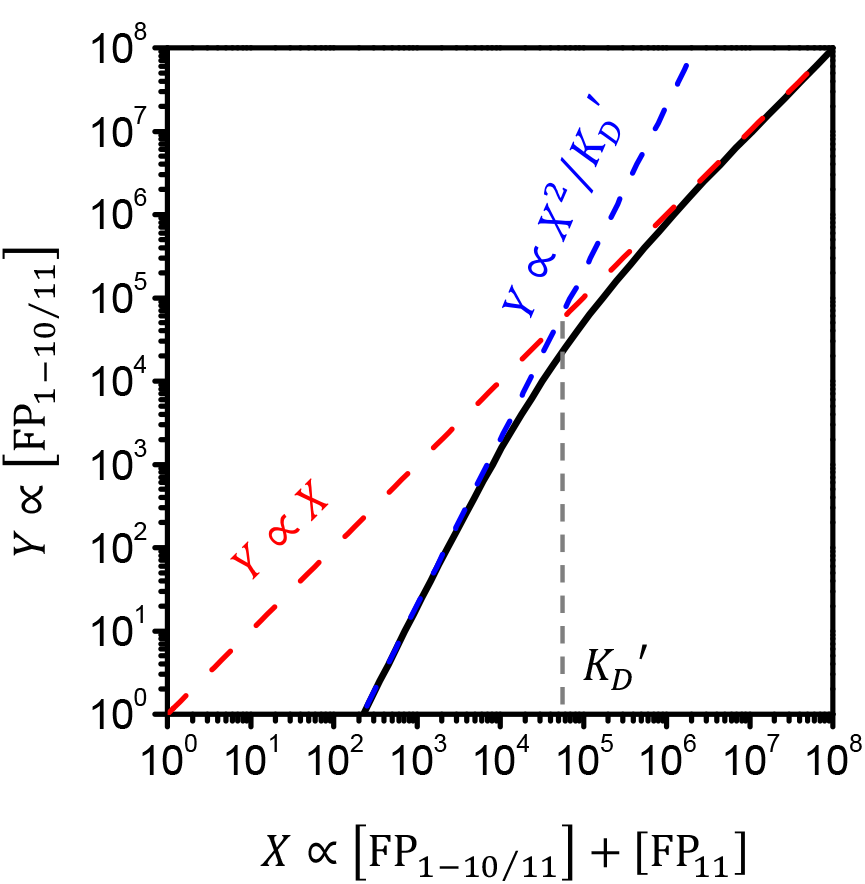

At a high fragment expression level or with a small *K_D_*′ (the case of GFP_1-10/11_), *X* is dominated by the first term of [FP_1–10/11_], making the right end of the curve approaching a slope of 1. On the other hand, at a low fragment expression level or with a large *K_D_*′ (the cases of mNG2_1-10/11_ and sfCherry2_1-10/11_), *X* is dominated by the second term of 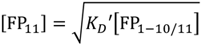, making the left end of the curve approaching a slope of 2.

In this latter regime, only a small fraction of the FP_11_ fragment is complemented with the FP_1-10_ fragment to reconstitute a functional fluorescent protein. Consequently, the apparent brightness of the FP_11_-labeled protein is lower than protein labeled by the full length FP. The complementation efficiency, defined as the fraction of FP_11_ complemented with FP_1-10_, is directly proportional to the concentration of FP_1-10_:

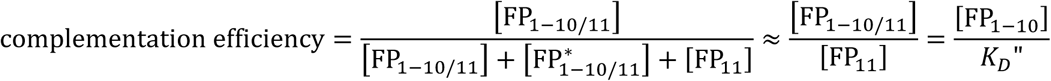

where *K_D_″* = *aK_D_″* (i.e. effective dissociation constant with respect to FP_1-10_ instead of FP_11_). This relationship holds true both for our FACS experiment and for any general applications of FP_11_ labeling. In other words, the complementation efficiency improves with higher expression level of the FP_1-10_ fragment. Therefore, overexpression of mNG2_1-10_ and sfCherry2_1-10_ improves the apparent brightness of the FP_11_ labeled proteins. This improvement saturates when the concentration of FP_1-10_ is approaching *K_D_″*. Therefore, a trade-off must be considered between the diminishing brightness improvement and the potential cytotoxicity of high FP_1-10_ expression.

Obviously, a better approach to improve the apparent brightness / complementation efficiency is to reduce *K_D_*′ (equivalent to reducing *K_D_″*). This is exactly what we tried to achieve by engineering sfCherry3C and sfCherry3V. In these cases, the improvement of *K_D_*′ can be measured by fitting the FACS data with a line at a slope of 2 and then measure the change in the intercept (fig. S4).

From sfCherry2 to sfCherry3C, the *K_D_*′ decreased by:

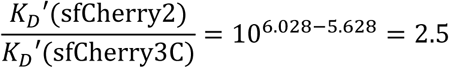

From sfCherry2 to sfCherry3V, the *K_D_*′ decreased by:

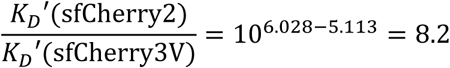

## Supplementary Tables

**Supplementary Table 1:**
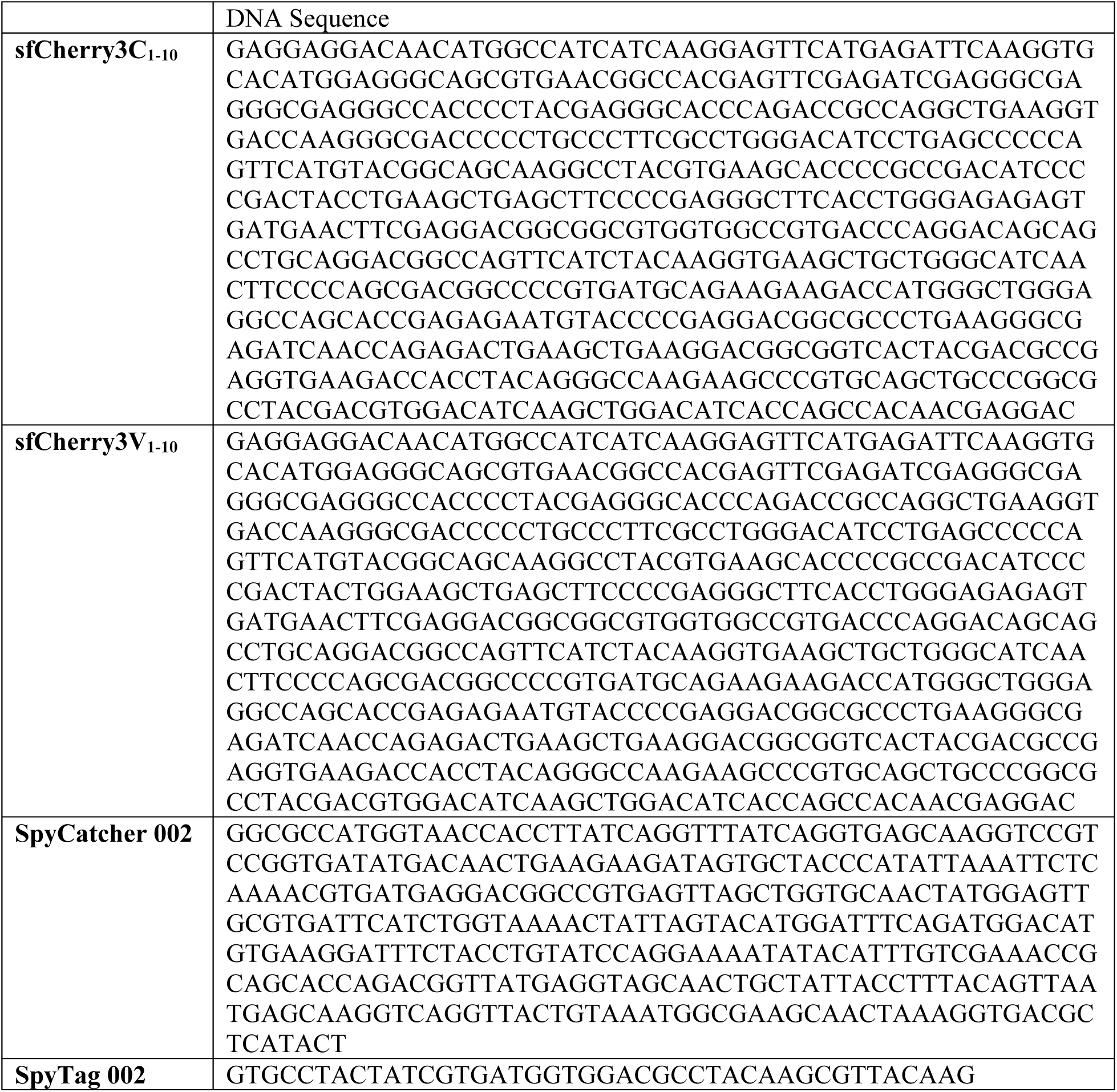
**Sequence of sfCherry3C_1-10_, sfCherry3V_1-10_, SpyCatcher 002 and SpyTag 002.**

**Supplementary Table 2:**
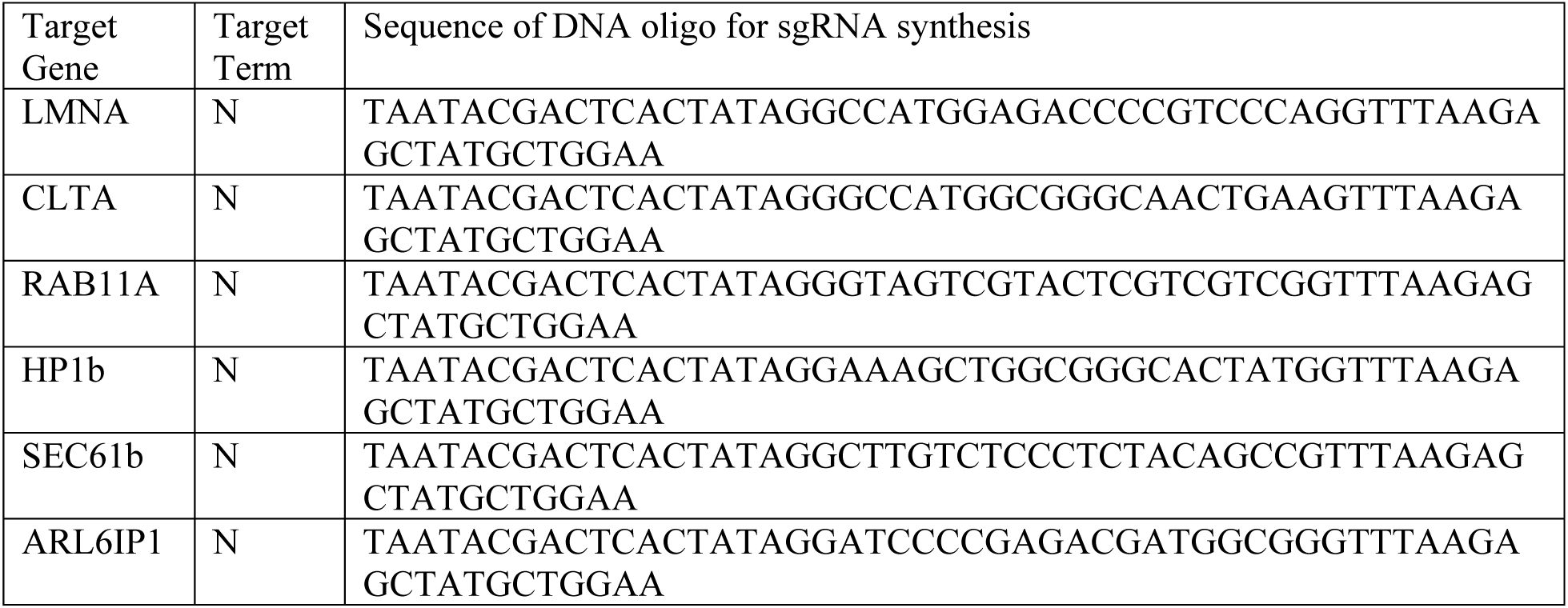
**DNA sequence used for sgRNA in vitro transcription**

**Supplementary Table 3:**
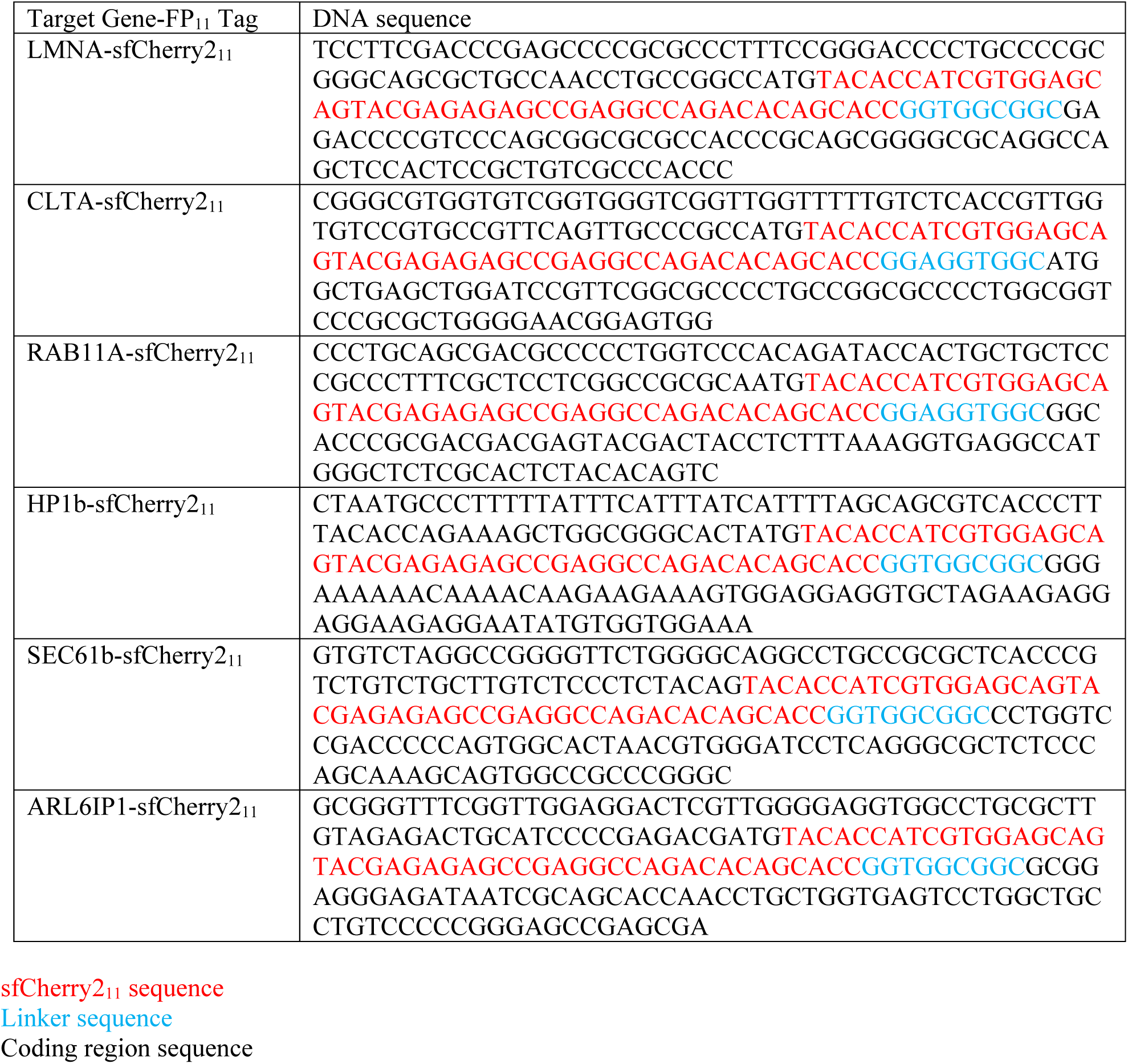
**Oligo-nucleotide donor DNA sequence**

